# The interpeduncular nucleus blunts the rewarding effect of nicotine

**DOI:** 10.1101/2023.10.05.561018

**Authors:** Joachim Jehl, Maria Ciscato, Eléonore Vicq, Gabrielle Dejean de la Batie, Nicolas Guyon, Sarah Mondoloni, Jacinthe Frangieh, Claire Nguyen, Jean-Pierre Hardelin, Fabio Marti, Pierre-Jean Corringer, Philippe Faure, Alexandre Mourot

**Affiliations:** Brain Plasticity Unit, CNRS, ESPCI Paris, PSL Research University, 75005 Paris, France; Sorbonne Université, Inserm, CNRS, Neuroscience Paris Seine - Institut de Biologie Paris Seine (NPS - IBPS), 75005 Paris, France; Institut Pasteur, Université de Paris, CNRS UMR 3571, Channel-Receptors Unit, Paris, France

## Abstract

Nicotine, by stimulating ventral tegmental area (VTA) dopaminergic neurons, has a rewarding effect that drives tobacco consumption. In turn, the interpeduncular nucleus (IPN) is thought to become activated at high nicotine doses to restrict drug intake. However, the dynamics of the IPN response to nicotine and its impact on the rewarding effect of the drug remain unknown. To address this issue, we have developed a genetically-modified mouse model, in which a “suicide” antagonist of nicotinic acetylcholine receptors (nAChRs) selectively attaches to a designer β4 nAChR subunit. By locally infusing this antagonist in the IPN, we achieved pharmacologically-specific and sustained antagonism of nAChRs containing the β4 subunit. By combining this chemogenetic method with *in vivo* electrophysiology, we show that even at low doses, nicotine activates and inhibits two different populations of IPN neurons, and that β4-containing nAChRs are only involved in the activation response. Furthermore, blocking the response to nicotine selectively in the IPN increased both the sensitivity of the VTA to the drug and its rewarding effect in a conditioned place preference paradigm. These findings indicate that the IPN is engaged across a large range of nicotine doses and acts as a regulatory brake on the nicotine reward circuit.

## Introduction

Nicotine is an addictive chemical substance found in the tobacco plant. Even though the percent of cigarette smokers currently decreases worldwide, there is a dramatic rise in the use of electronic nicotine delivery systems, particularly in teenagers.^1^ Understanding the physiological mechanism of nicotine addiction is thus crucial for developing effective strategies to help individuals break free from its grip. Nicotine exerts its effects by binding to nicotinic acetylcholine receptors (nAChRs). There are nine different alpha (α2-9) and three different beta (β2-4) nAChR subunits in the brain, which assemble to form a variety of hetero- and homo- pentameric structures.^2^

The initiation of addiction to nicotine involves the mesolimbic dopamine (DA) reward system.^3^ Nicotine acts as a positive reinforcer by activating α4β2 nAChRs located in the ventral tegmental area (VTA),^4–8^ which leads to the release of DA in the nucleus accumbens.^9^ Nicotine also elicits negative effects, depending on the dose. In particular, a reinforcing dose that motivates drug intake^5^ also drives anxiety.^9^ At higher doses, nicotine has been described as aversive, thus motivating avoidance.^10–13^ These competing reinforcing and aversive properties of nicotine, together with the feeling of satiety, are consistent with the observation that humans and laboratory rodents titrate their nicotine intake when high doses are available.^5,11,12,14^

The aversive effect of high-dose nicotine involves the neural pathway projecting from the medial habenula (Mhb) to the interpeduncular nucleus (IPN).^5,10,11^ Neurons belonging to this pathway express high densities of rare nAChR subunits, notably α3, α5 and β4.^15–17^ In particular, the α3 and β4 subunits are almost absent in the VTA^18^ and in other parts of the brain.^17,19^ Knock-out mice lacking α5^11^ or β4^12,20,21^ consume more nicotine than wild-type (WT) mice, especially at high doses, and their consumption level is restored upon viral re-expression of these nAChR subunits in the Mhb^11,20^ or IPN.^12,20,21^ Similarly, knocking down α3 selectively in the Mhb or IPN^22^, or pharmacologically blocking α3β4 nAChRs in the IPN^23^, results in increased self-administration of nicotine in the rat. Because the α3β4 nAChR has lower affinity for nicotine than the α4β2 nAChR^2,24^, one current hypothesis is that nicotine is rewarding at low doses because it activates primarily α4β2 receptors of the VTA, while it is aversive at high doses because only then does it activate α3β4 nAChRs of the MHb-IPN axis.^5,13^ It is currently unknown whether the IPN also responds to low doses of nicotine *in vivo*, and how this may affect the balance between the reinforcing and aversive properties of the drug.

## Results

### Conditioning at a low dose of nicotine in β4^-/-^ mice

To assess the rewarding properties of low-dose nicotine, we employed an unbiased conditioned place preference (CPP) paradigm. Acute intraperitoneal administration of nicotine at a dose of 0.5 mg/kg consistently induces a preference for the drug-paired chamber in C57BL/6 mice.^68,25–27^ However, lower doses of nicotine typically produce neither aversion nor preference for the drug in WT mice.^27–29^ Accordingly, a low dose of nicotine (0.2 mg/kg) did not induce a preference compared to saline in wild-type mice, but did induce nicotine preference in mutant mice lacking the nAChR β4 subunit (Figure 1A). This subunit is abundant in the MHb-IPN pathway but virtually absent in other brain regions including the VTA,^18^ suggesting that low doses of nicotine may act through β4-containing nAChRs, conceivably within the MHb-IPN pathway, to modulate the rewarding effect of the drug. To address this question, we devised a method that enables long-lasting and exclusive manipulation of β4-containing nAChR subtypes in targeted brain regions.

**Figure 1:**
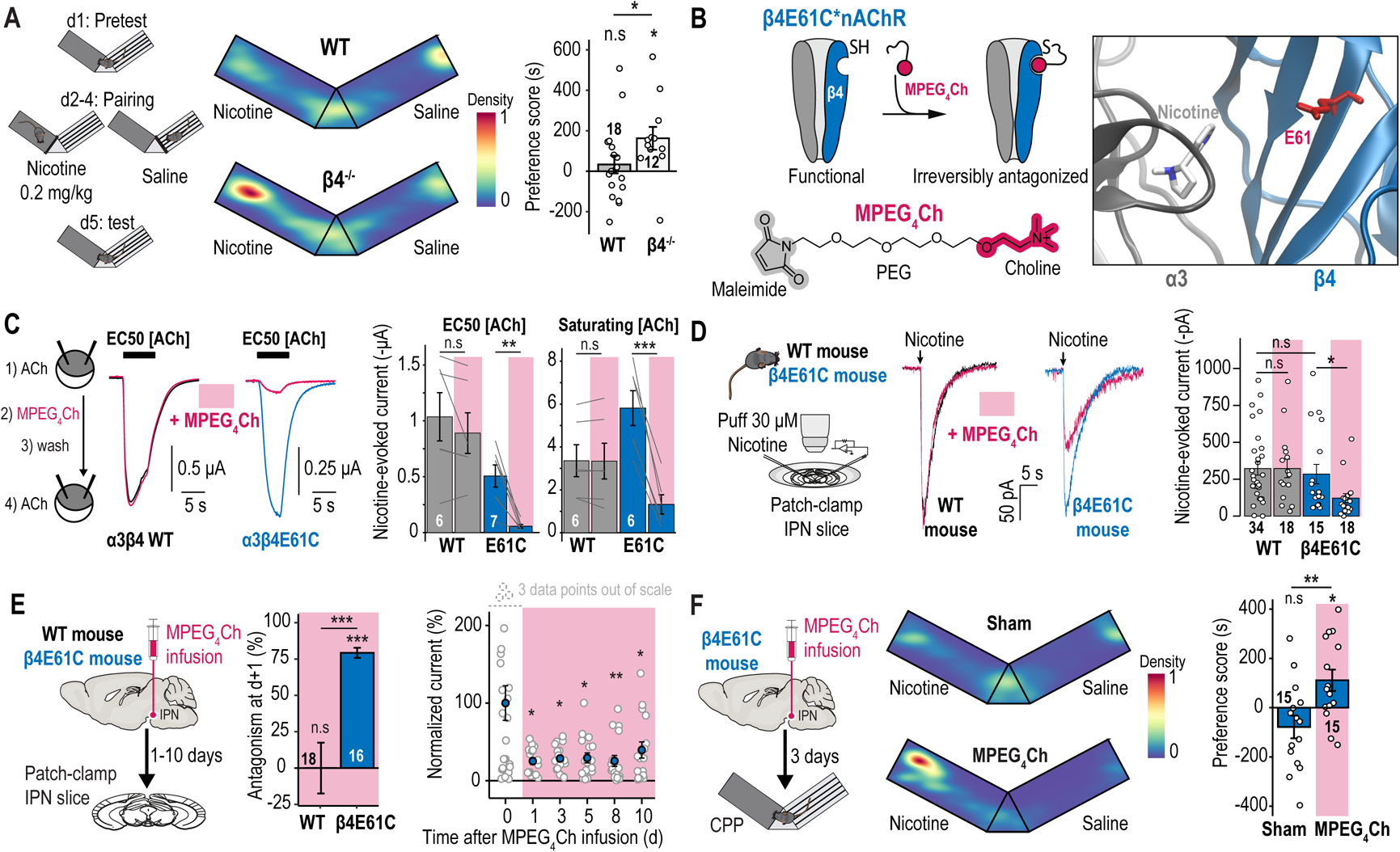
Chemogenetic method for sustained antagonism of β4-containing nAChRs. **A.** Left, unbiased CPP protocol. Middle, representative heat maps for a WT and a β4^-/-^ mouse. Right, preference for the nicotine-paired chamber. **B.** Top left, β4E61C-containing nAChR irreversibly antagonized after covalent attachment of MPEG_4_Ch. Bottom left, chemical structure of MPEG_4_Ch. Right, side view of the agonist binding site in the α3β4 nAChR in complex with nicotine (PDB: 6pv7)^44^, with E61 shown in red. **C.** Left, ACh-induced currents recorded using two-electrode voltage clamp, before and after MPEG_4_Ch treatment. Right, average currents before and after MPEG_4_Ch treatment. **D.** Left, patch-clamp recording with puff application of nicotine on IPN slices. Representative (middle) and average (right) nicotine-induced currents recorded from slices treated or not with MPEG_4_Ch. **E.** Left, local infusion of MPEG_4_Ch in the IPN, 1 to 10 days before preparing the slices. Middle, percent inhibition of nicotinic current in IPN neurons one day after MPEG_4_Ch infusion. Right, nicotine-induced current 1 to 10 days after MPEG_4_Ch infusion. **F.** Left, nicotine CPP protocol after MPEG_4_Ch treatment. Middle, representative animal positions. Right, preference for the nicotine-paired chamber.

### Designing chemogenetic tools for sustained nAChR antagonism

We have previously developed a chemogenetic method to manipulate specific nAChR subtypes in the living mouse with light.^6,30^ It relies on the covalent attachment of a photoswitchable tethered ligand called MAHoCh near the acetylcholine binding pocket, by replacing the amino acid glutamate at position 61 by a cysteine (E61C) in the β2 or β4 nAChR subunit. While this method allows rapid and reversible control of nicotinic signaling, its *in vivo* application requires mice to be continuously connected to an external light source, which poses challenges for long-term behavioral experiments.^31^ To overcome this limitation, we sought to extend the chemogenetic toolbox for nAChR control, by developing a method that would allow sustained receptor antagonism without the need for light stimulation. The idea was to covalently attach a synthetic antagonist to a genetically-encoded cysteine tag at the nAChR surface, in close proximity to the agonist binding pocket (Figure 1B). The antagonist acts as a suicide inhibitor: its covalent attachment to the cysteine residue increases the local concentration of the antagonist in the vicinity of the receptor and leads to permanent occupation of the agonist binding pocket, resulting in sustained antagonism. The newly developed tethered antagonist, named MPEG_4_Ch, contains a cysteine-reactive maleimide (M) group at one end, a choline (Ch) ligand at the other end, and a flexible poly-ethylene glycol (PEG) linker in between (Figure 1B). To maximize the versatility of our chemogenetic toolbox, we designed MPEG_4_Ch so that it antagonizes nAChRs after attachment to the very cysteine residue (E61C) that makes nAChRs photocontrolable (Figure 1B).^6,30^ The size of the PEG linker (consisting of four ethylene glycol units) was thus chosen to match that of the photoswitchable azobenzene linker in MAHoCh (Figure S1A).

To experimentally validate our strategy, we heterologously expressed WT and cysteine-substituted α3β4 nAChRs in Xenopus oocytes. The mutation E61C in β4 had minimal effect on the receptor sensitivity to acetylcholine (Figure S1B), indicating that the mutant receptor is functional, which is consistent with our previous findings.^30^ After incubation of the oocytes with 15 μM MPEG_4_Ch and subsequent washing, there was a robust inhibition of α3β4E61C receptors at both EC_50_ and saturating concentrations of acetylcholine (∼90% and 80%, respectively, Figure 1C), but no effect on WT receptors, indicating that the cysteine substitution is required for the antagonistic action of MPEG_4_Ch.

### Knock-in mice for the manipulation of endogenous nAChRs

To manipulate nicotinic signaling while preserving endogenous nAChR expression patterns, we produced a knock-in mouse line carrying a single point mutation at codon position 61 in the gene encoding the nAChR β4 subunit (β4E61C, Figure S1C). The amplitudes of nicotine-induced currents were identical in the knock-in and the WT mice (Figure 1D), suggesting unchanged nAChR expression profiles in the neurons of mutant mice, as expected. We also verified that IPN neurons of these mice had no obvious electrophysiological alteration (Figure S1D-G), and that the mice had no discernable phenotypical alteration (Figure S1H-I). These results indicate that the β4E61C minimally (if at all) perturbs nicotinic signaling.

We then assessed the ability of MPEG_4_Ch to inhibit β4-containing nAChRs in neurons of the knock-in mouse. IPN brain slices of adult WT and β4E61C mice were treated with MPEG_4_Ch, extensively washed, and the amplitude of nicotine-induced currents was measured in whole-cell recordings. MPEG_4_Ch treatment markedly decreased the amplitude of the response to nicotine in β4E61C mice (> 60%), but had no effect in WT mice (Figure 1D). This result confirms that β4-containing nAChRs are the main receptor subtypes in the IPN,^12^ and that the engineered E61C mutation is required for MPEG_4_Ch-induced nAChR antagonism.

Next, we examined the ability of MPEG_4_Ch to conjugate to β4E61C-containing nAChRs *in vivo*. MPEG_4_Ch was infused into the IPN of anesthetized mice, and brain slices were prepared on the following day. MPEG_4_Ch reduced nicotine-induced currents in IPN neurons of β4E61C mice by about 80%, but had no effect in WT mice (Figure 1E). Since MPEG_4_Ch irreversibly reacts with cysteine residues, the antagonism should persist as long as the MPEG_4_Ch-conjugated receptors remain at the cell surface. To investigate the duration of antagonism, brain slices were prepared 1 to 10 days after a single injection of MPEG_4_Ch in the IPN, and we found that the antagonistic effect persisted up to 10 days (Figure 1E). Therefore, MPEG_4_Ch can be used as a cysteine-reactive covalent ligand, providing efficient, long-lasting, and subtype-specific antagonism of β4E61C-containing nAChRs.

### Blocking the IPN is sufficient to induce a preference to low-dose nicotine

We then investigated whether the rewarding effect of low-dose nicotine in β4 knock-out mice (Figure 1A) was due to the absence of β4 in the IPN. We selectively blocked β4 nAChRs in the IPN with MPEG_4_Ch, and carried out the CPP test. Sham-injected β4E61C mice served as the control group. MPEG_4_Ch was injected 3 days prior to the conditioning phase, to ensure continuous nAChR antagonism throughout the protocol. Only the MPEG_4_Ch-treated mice, but not the sham-treated mice, developed a preference for low-dose nicotine (Figure 1F), therefore reproducing the result with the knock-out mice, and indicating that β4-containing nAChRs of the IPN play a key role in setting the rewarding value of nicotine, especially at low dose.

### Dose-dependent effects of nicotine on IPN neurons *in vivo*

To understand the mechanism by which the IPN affects the rewarding effect of the drug, we first assessed the sensitivity of the IPN to acute nicotine injections *in vivo*. We used a microdrive multielectrode manipulator to record the electrical activity of IPN neurons in anesthetized mice (Figure 2A). Acute intravenous (i.v.) injection of 30 μg/kg nicotine, a reinforcing dose typically used to probe nicotine response in VTA neurons^6,9,32^, evoked opposing responses (activation *vs.* inhibition) in IPN neurons (Figure 2A). Of the 168 neurons recorded, 97 (58 %) showed an increase in firing rate, while 71 (42 %) showed a decrease, and these variations were absent with saline injection (Figure 2B-C and S2A). The two populations of neurons could not be distinguished based on their spontaneous firing rates (Figure S2B), but there was a correlation between the spontaneous firing rate and the amplitude of the response to nicotine, in both activated and inhibited cells (Figure S2C). These results are consistent with previous data obtained using the juxtacellular technique in neurons that were labelled and confirmed to be located in the IPN.^12^

**Figure 2:**
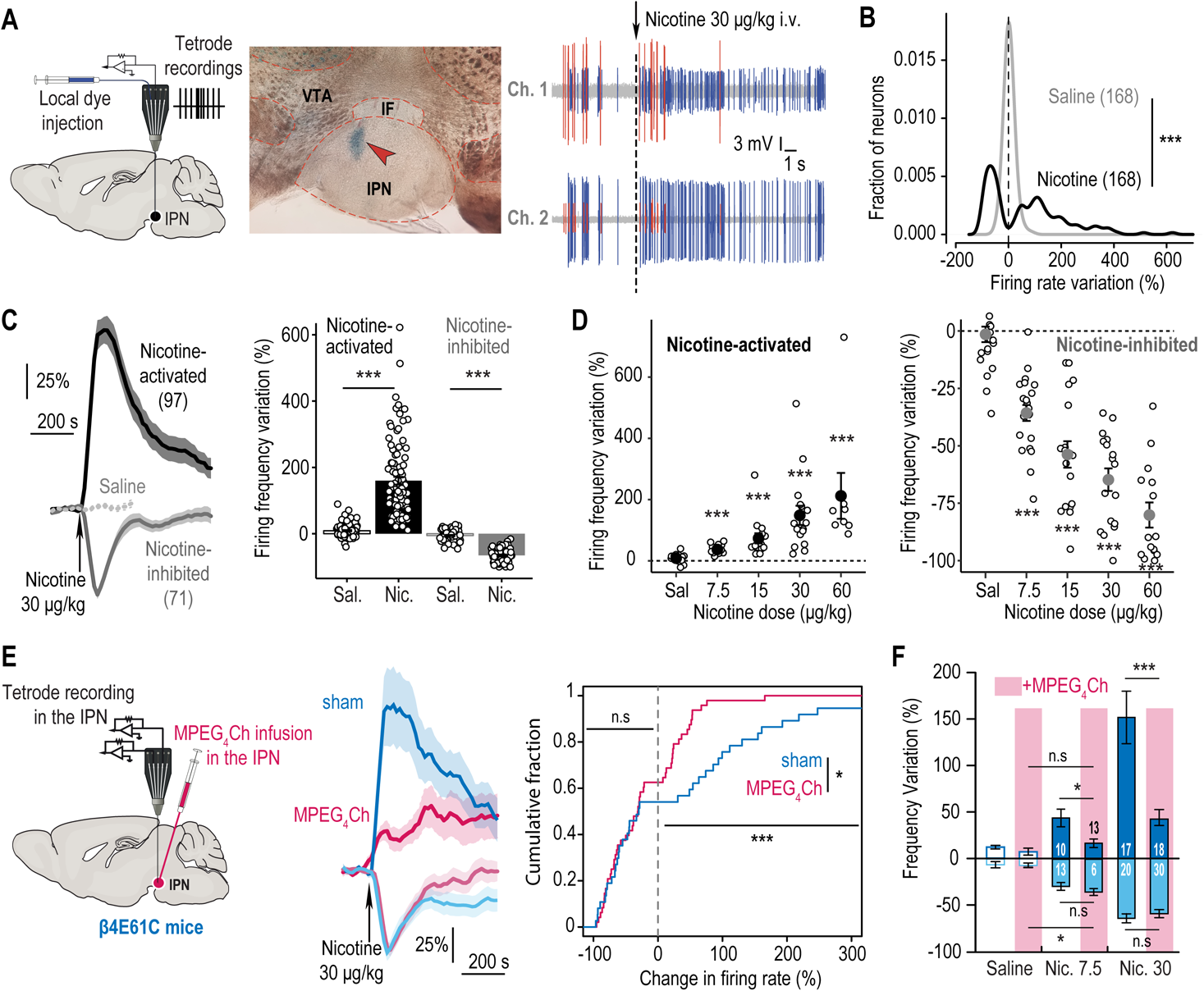
β4-containing nAChRs are specifically involved in the activation of IPN neurons by nicotine. **A.** Left, schematics of the microdrive multielectrode manipulator setup for tetrode recordings. Middle, a blue dye was injected at the recording site coordinates. Right, two IPN neurons recorded simultaneously on two channels, one activated (blue) and one inhibited (red) by nicotine. **B.** Distribution of the responses to i.v. injections of nicotine or saline. **C.** Time course (left) and firing frequency variation (right) after i.v. nicotine (or saline) injections, for nicotine-activated and -inhibited IPN neurons. **D.** Variation in firing frequency for nicotine-activated and -inhibited IPN neurons, following i.v. injections of different doses of nicotine. Neurons were classified as activated or inhibited based on their response at the 30 μg/kg dose (or alternatively 60 μg/kg for a few neurons not recorded at 30 μg/kg). **E.** Left, infusion of MPEG_4_Ch (or saline for controls) in the IPN of β4E61C mice, followed by tetrode recordings in the IPN. Middle, time course of responses to nicotine injection. Right, cumulative fraction of the percent change in firing rate in response to nicotine injection. **F.** Maximum amplitude of activation or inhibition following an i.v. injection of nicotine or saline.

We then assessed the sensitivity of IPN neurons to a range of nicotine doses (7.5 to 60 μg/kg) and found that the effect of nicotine was dose-dependent, but that the polarity of the response (*i.e.*, activation or inhibition) was not. Indeed, neurons that were activated by nicotine at 30 μg/kg were activated across all doses tested, and similarly, inhibited neurons maintained their inhibition at all doses (Figure 2D). The two populations of neurons are thus distinct and not overlapping. Most of the neurons (36/37), whether activated or inhibited, responded to the lowest dose of 7.5 μg/kg.

### β4-containing nAChRs mediate activation of IPN neurons by nicotine

We employed our new chemogenetic strategy to assess the contribution of β4-containing nAChRs to the response of IPN neurons to nicotine *in vivo*. MPEG_4_Ch (or saline for the control sham group) was locally infused in the IPN of β4E61C mice with the microdrive multielectrode manipulator, and the electrical activity of IPN neurons was recorded after a couple of hours (Figure 2E). MPEG_4_Ch did not affect the spontaneous electrical activity of IPN cells (Figure S2E). However, the amplitude of nicotine-induced activation was markedly reduced in MPEG_4_Ch-treated animals, for both 7.5 and 30 μg/kg doses (Figure 2E-F). In contrast, the amplitude of nicotine-induced inhibition was unaffected by the MPEG_4_Ch treatment, in agreement with observations previously made in β4^-/-^ mice^12^. We reproduced these experiments in WT mice and showed that MPEG_4_Ch, without the ability to anchor to nAChRs, had no effect on the response to nicotine, confirming the molecular specificity of the method (Figure S2D). Together, these results indicate that IPN β4-containing nAChRs are recruited even at low nicotine doses, and predominantly contribute to the nicotine response in activated neurons.

### The IPN is more sensitive to nicotine than the VTA

Our results indicate that IPN neurons respond to i.v. nicotine doses as low as 7.5 μg/kg. In comparison, previous studies have reported that VTA DA neurons start responding at twice this dose (15 μg/kg).^8,31,32^ To compare the relative sensitivities of the VTA and IPN to nicotine, we conducted two independent experiments. First, we recorded bulk neuronal activity in each brain region using fiber photometry. The VTA or IPN of WT mice were transduced with the calcium sensor GCaMP7c, and proper transduction and fiber location were verified with *post-hoc* immunochemistry (Figures 3A-B). Mice were anesthetized and different doses of nicotine (7.5 - 60 μg/kg) were injected i.v. while monitoring the GCaMP signal. Nicotine increased the calcium signal in the IPN across all doses tested (Figure 3A), while responses in the VTA were only detected from the 30 μg/kg dose onward (Figure 3B).

**Figure 3:**
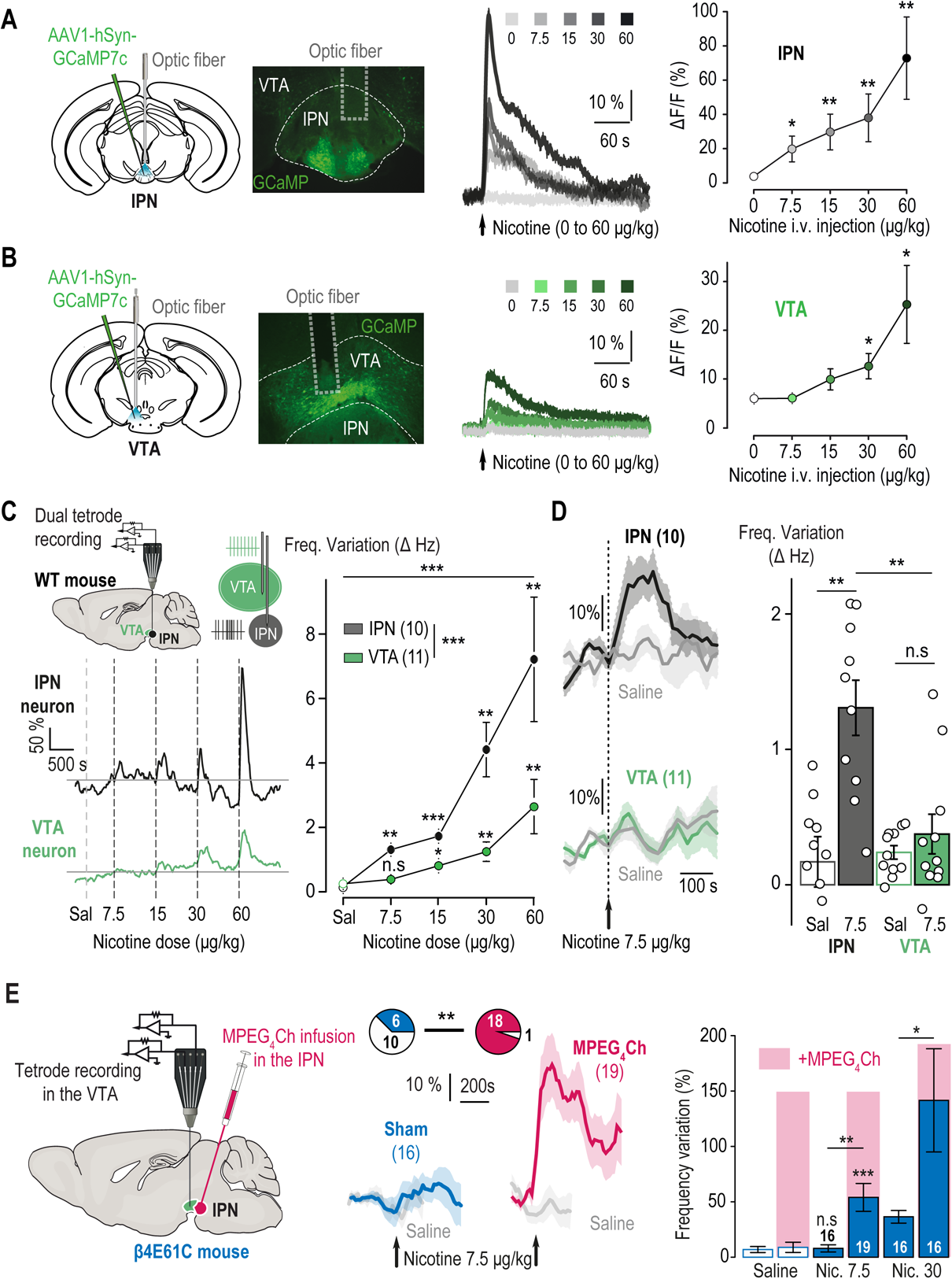
The IPN decreases the VTA response to nicotine. **A.** Transduction of the IPN with AAV1-hSyn-GCaMP7c, followed by implantation of the optic fiber, and a representative coronal brain slice showing the fiber tip and proper expression of GCaMP7c. Time course (middle) and maximum amplitude (right) of the GCaMP7c response after i.v. injection of nicotine at different doses. **B.** Same as in A) but for the VTA. **C.** Top left, simultaneous tetrode recordings with one electrode in the IPN and another one in the VTA. Representative (bottom left) and average (right) change in firing frequency in IPN and pDA neurons following i.v. injection of nicotine. **D.** Time course (left) and average change (right) of the response of IPN and VTA neurons to an i.v. injection of 7.5 μg/kg nicotine or saline. **E.** Left, tetrode recordings in the VTA following local infusion of MPEG_4_Ch into the IPN of a β4E61C mouse. Middle, time course of the response to an i.v. injection of 7.5 μg/kg nicotine (or saline). Only about a third (6/16) of the pDA neurons responded to the nicotine injection in sham-treated mice, whereas almost all of them (18/19) responded in MPEG_4_Ch-treated mice. Right, average response to an i.v. injection of nicotine or saline.

In a second step, we positioned one tetrode in the VTA and another one in the IPN to simultaneously record in the two structures. Overall, the amplitude of firing frequency variation was greater in IPN neurons than in putative DA (pDA) neurons of the VTA at all nicotine doses tested (Figure 3C). VTA pDA neurons started to respond at a dose of 15 μg/kg (Figure 3C), in agreement with previous recordings of identified VTA dopaminergic neurons,^7,9^ while IPN neurons responded at all doses. Specifically, the lowest nicotine dose tested (7.5 μg/kg) increased the firing rate of IPN neurons but not of VTA pDA neurons (Figure 3D). Collectively, these optical and electrophysiological recordings strongly suggest that IPN neurons respond to nicotine with greater sensitivity and greater amplitude than VTA DA neurons.

### The IPN lowers the rewarding effect of nicotine

We then hypothesized that the response to nicotine in the IPN impacts on the response in the VTA, potentially explaining why animals with a reduced nicotine response in the IPN showed an increased preference for nicotine. We took advantage of our newly-developed chemogenetic approach to inhibit the response to nicotine solely in the IPN while recording in the VTA. MPEG_4_Ch (or saline for control mice) was infused into the IPN of β4E61C mice, and the response of VTA neurons to nicotine i.v. injections was subsequently measured with tetrodes. Strikingly, VTA neurons of MPEG_4_Ch-treated mice, but not of control mice, responded to the injection of low-dose nicotine (7.5 μg/kg, Figure 3E). Furthermore, the amplitude of the response to nicotine was greater for MPEG_4_Ch-treated mice than for control animals, even when a higher, more standard dose of 30 μg/kg was administered. To rule out the possibility that MPEG_4_Ch may have diffused outside the IPN, we carried out two control experiments. First, we infused MPEG_4_Ch in the VTA of β4E61C mice, and found that this treatment had no effect on the response of VTA pDA neurons to nicotine (Figure S3A), in agreement with the reported absence of β4-containing nAChRs in the VTA.^18^ Then, we checked whether infusion of MPEG_4_Ch in the IPN could affect neurons of the MHb, a distant brain region with dense expression of β4-containing nAChRs. Incubation of MHb slices with MPEG_4_Ch profoundly decreased the amplitude of nAChR currents, confirming the strong expression level of β4 in the MHb,^17^ but infusion of MPEG_4_Ch in the IPN had no effect on the response of MHb neurons to nicotine (Figure S3B). Taken together, these experiments indicate that the effect of MPEG_4_Ch on VTA neurons is unlikely to be attributed to the its diffusion outside the IPN. Therefore, blocking the response to nicotine selectively in the IPN boosted the amplitude of the response to the drug in the VTA, providing a mechanistic understanding of the increased sensitivity to nicotine in β4^-/-^ mice.^21^

## Discussion

Making receptors light-controllable is a valuable method for probing neurotransmission with high spatiotemporal precision.^31,33^ However, it is not convenient for manipulations that require long-term effects, particularly in behaving mice. To overcome this limitation, we expanded the toolbox for controlling nAChRs, by developing a knock-in mouse line (β4E61C) and a corresponding suicide inhibitor (MPEG_4_Ch) that forms a covalent bond with the cysteine amino acid installed in the knock-in mouse. The method provides efficient antagonism (approximately 80%) with immediate onset (maleimides react within minutes with thiols^34^), sustained effect (several days), absolute pharmacologically specificity (nAChRs containing the β4 subunit only) and anatomical precision. This combination of properties is difficult to achieve with traditional pharmacological tools, because small molecules rarely show receptor-subtype specificity, and their effects ware off rapidly due to diffusion in brain tissues. While gene knock-down strategies using shRNAs can achieve long-lasting inhibition of receptors in targeted tissue, the overall degree of silencing was found to vary tremendously,^35,36^ and the effect takes weeks to occur.

In this study, we used this chemogenetic method in knock-in mice, where the WT nAChR β4 subunit has been replaced with a “silent” mutant subunit, ensuring proper expression profiles and untouched nicotinic transmission. We provide here direct evidence that β4- containing nAChRs are functionally expressed at high levels in both IPN and MHb neurons. In principle, this mouse line could also be used in combination with previously-developed photoswitchable tethered ligands^6,30^ to afford rapid, optical manipulation of endogenous nAChRs. Alternatively, local injection of a viral vector could be used to get restricted expression of the β4_E61C_ subunit in targeted cell types.^6,33^ In β4E61C knock-in mice, a single bolus dose of MPEG_4_Ch resulted in robust receptor inhibition for several days, which suggests that bioconjugation of MPEG_4_Ch to the thiol group of cysteine residues is fully irreversible, as demonstrated for other PEG-substituted maleimide reagents.^34^ This finding also suggests that the lifetime of nAChRs *in vivo* is in the order of weeks, which is compatible with the median lifetime of about 10 days measured for neuronal receptors in the mouse brain.^37^

A previous study utilizing c-fos imaging reported that the IPN only responds to high, aversive doses of nicotine, and not to a lower, rewarding dose.^11^ This led to the hypothesis that the IPN is only engaged at high nicotine doses to limit drug intake.^5,13^ By recording IPN neurons in live animals, we have revealed their sensitivity to a wider range of nicotine concentrations. We found that the IPN responds to concentrations of nicotine that do not activate the VTA. More importantly, we found that the effect of low-dose nicotine on the IPN translates into a reduced potency of the drug in the VTA. This implies that the mesolimbic DA and MHb-IPN pathways, which encompass the IPN and VTA, do not work independently. Our results also uncover that low, sub-rewarding doses of nicotine can act through α3β4 nAChRs, which are commonly considered as low-affinity receptors.^24^ We suggest that the IPN acts as a brake on the VTA to decrease the rewarding value of the drug and ultimately to limit drug intake.

The mechanism by which the IPN reduces the effect of nicotine in the VTA remains unclear. Currently, there is no evidence for a direct neural connection from the IPN to the VTA.^38,39^ However, the laterodorsal tegmentum (LDTg) has been identified as an important functional relay between these two structures. GABAergic projections from the IPN to the LTDg are modulated by nicotine, and their optogenetic stimulation elicits avoidance behavior.^13^ The LDTg, in turn, sends excitatory glutamatergic and cholinergic inputs to the VTA, and their activation mediates reward.^40,41^ It is thus plausible that nicotine, by activating GABAergic projections from the IPN to the LDTg, gates afferent excitatory signals onto the VTA, thereby decreasing the direct impact of the drug on DA neurons. The IPN is nevertheless a complex, heterogenous structure with multiple subnuclei and different populations of GABAergic projection neurons and interneurons, each with precise molecular and anatomical signatures.^38,42,43^ Nicotine-activated and -inhibited neurons could not be identified solely on basis of electrophysiological criteria. Further investigations are necessary to determine whether IPN neurons projecting to the LDTg are activated or inhibited by nicotine, and whether the IPN regulates the encoding of rewards other than nicotine in the VTA.

## Acknowledgements

The authors would like to thank Romain Durand-de Cuttoli (Sorbonne University, Paris, France) for his help with electrophysiology experiments, and the animal facilities at IBPS Sorbonne University and ESPCI Paris.

## Author contribution

Conceptualization: AM and PF

Methodology: AM

Validation: FM, PJC, PF, AM

Formal analysis: JJ, EV, MC, GDB, JF, NG, SM, CN, AM

Investigation: JJ, EV, MC, GDB, JF, NG, SM, CN

Writing, original draft: AM, JJ, MC, JPH

Writing, review and editing: all authors

Visualization: JJ, MC, EV, AM

Supervision: FM, PJC, PF, AM

Funding acquisition: PJC, PF, AM

## STAR Methods

### Animals

Mice were housed at 18-23 °C under a 12-hour light/dark cycle, with food and water available ad libitum. WT mice were on the C57BL/6 N genetic background. Female and male mice were used for the electrophysiological recordings, whereas only male mice were used for the fiber photometry and behavioral experiments. All behavioral tests were conducted during the light period of the animal cycle. All procedures for animal maintenance, surgery and behavioral tests used protocols that were in accordance with the recommendations for animal experiments issued by the European Commission directives 2010/63, and approved by Sorbonne University and Paris Sciences et Lettres (PSL) University.

The transgenic mouse line Chrnb4E61C was produced on a C57BL/6 N genetic background using homologous recombination, at the core facility Phenomin (Institut Clinique de la Souris, Illkirch, France). Exons 3 and 4 were fused in both lines to prevent alternative splicing, owing to the overlap between the codon for E61 and the splice acceptor site. The following primers were used to genotype Chrnb4E61C mice: forward 5’-CCAAGAGAGATTTAGGTGACAGCTGG-3’ and reverse 5’-CCTAACTCTAGACTTTGGCCACCCC-3’. Mice were bred as homozygotes.

### Chemicals

MPEG_4_Ch (maleimide - tetraethylene glycol - choline) chloride (purity 99.8%) was custom-synthetized by Broadpharm (San Diego, CA) and stored at −20°C in water-free DMSO. The nicotine used for all experiments is a nicotine hydrogen tartrate salt (Sigma-Aldrich, USA). For tetrode and fiber photometry recordings, we performed i.v. injection of both nicotine (example dose of 30 μg/kg) and saline (water with 0.9% NaCl) solutions. For the conditioned place preference (CPP) paradigm, mice were injected intra-peritoneally (i.p.) with nicotine at 0.2 mg/kg and immediately placed in the apparatus. All solutions were prepared in the laboratory.

### Two-electrode voltage-clamp

*Xenopus laevis* oocytes were obtained from EcoCyte Bioscience, Germany and and from Tefor Paris-Saclay UAR2010 and maintained in modified Barth’s medium (87.34 mM NaCl, 1 mM KCl, 0.66mM NaNO_3_, 0.75 mM CaCl_2_, 0.82mM MgSO_4_, 2.4 mM NaHCO_3_, 10 mM HEPES pH 7.6). Defolliculated oocytes were submitted to intranuclear injection of ∼2-6 ng of cDNA and kept at 18 °C for 2-3 days before recording.

Recordings were performed with a Digidata 1550A digitizer (Molecular Devices), an Axon Instruments GeneClamp 500 amplifier (Molecular Devices), an automated voltage-controlled perfusion system which controls an 8-port and a 12-port electric rotary valve (Bio-Chem Fluidics) both connected to a 2-way 4-port electric rotary valve (Bio-Chem Fluidics) and the pClamp 10.6 software (Molecular Devices).

Electrode pipettes were pulled using a Narishige PC-10 puller. Oocytes were perfused with Ringer’s buffer (100 mM NaCl, 2.5 mM KCl, 10 mM HEPES, 2 mM CaCl_2_, 1 mM MgCl_2_, pH 7.3). MPEG_4_Ch solutions were applied after dilution in Ringer’s buffer and all currents were measured at a holding potential of −60mV.

For all the experiments, the oocytes were perfused with Ringer’s buffer for 30 s, then 5 s with acetylcholine (concentrations as indicated in the figures) followed by a 1 to 2-minute wash. For the experiments with MPEG_4_Ch, the oocytes were first perfused with acetylcholine followed by a 1 to 2-minute wash, then treated with a perfusion of 5 min with MPEG_4_Ch (15 μM in Ringer’s), followed by another 1-2 minute wash and a 5 s acetylcholine perfusion. Acetylcholine responses were measured at the peak of the response.

### Patch-clamp electrophysiology

Mice were deeply anesthetized with an i.p. injection of a mixture of ketamine (150 mg/kg, Imalgene 1000, Merial) and xylazine (60 mg/kg, Rompun 2%, Bayer). Coronal midbrain sections (250 μm) were sliced using a Compresstome (VF-200; Precisionary Instruments) after intra-cardiac perfusion of cold (0-4°C) sucrose-based artificial cerebrospinal fluid (aCSF) containing (in mM): 125 NaCl, 2.5 KCl, 1.25 NaH_2_PO_4_, 5.9 MgCl_2_, 26 NaHCO_3_, 25 sucrose, 2.5 glucose, 1 kynurenate. After 10 min at 35 C° for recovery, slices were transferred into oxygenated (95% CO_2_/ 5% O_2_) aCSF containing (in mM): 125 NaCl, 2.5 KCl, 1.25 NaH_2_PO_4_, 2 CaCl_2_, 1 MgCl_2_, 26 NaHCO_3_, 15 sucrose, 10 glucose at room temperature for the rest of the day. Slices were labeled individually with MPEG_4_Ch (50 μM) in oxygenated aCSF for 10 min, and transferred to a recording chamber continuously perfused at 2 ml/min with oxygenated aCSF.

Patch pipettes (5-8 MΩ) were pulled from thin-wall borosilicate glass (G150TF-3, Warner Instruments) using a micropipette puller (P-87, Sutter Instruments) and filled with a K-gluconate based intra-pipette solution containing (in mM): 135 KGlu, 10 HEPES, 0.1 EGTA, 5 KCl, 2 MgCl_2_, 2 ATP-Mg, 0.2 GTP-Na and 2 mg/mL biocytin (pH adjusted to 7.35). Cells were visualized using an upright microscope with a Dodt contrast lens and illuminated with a white light source (Scientifica). Whole-cell recordings were performed using a patch-clamp amplifier (Axoclamp 200B, Molecular Devices) connected to a Digidata (1550 LowNoise acquisition system, Molecular Devices). Currents were recorded in voltage-clamp mode at −60 mV. Signals were low-pass filtered (Bessel, 2 kHz) and collected at 10 kHz using the data acquisition software pClamp 10.5 (Molecular Devices). Electrophysiological recordings were extracted using Clampfit (Molecular Devices) and analyzed with R. To record nicotinic currents from IPN neurons, local puffs (200 ms) of nicotine tartrate (30 μM in aCSF, Sigma-Aldrich) were applied every minute, using a glass pipette (2-3 μm diameter) positioned 20 to 30 μm away from the soma and connected to a picospritzer (World Precision Instruments, adjusted to ∼2 psi).

### Stereotaxic surgeries

For virus and MPEG_4_Ch injections, as well as for optic fiber implantations, mice were exposed to a gas mixture of oxygen (1 L/min) and 3% isoflurane (Piramal Healthcare, UK) for induction of anesthesia, and then placed in a stereotaxic frame (David Kopf Instruments, CA, USA) under 1% isoflurane to maintain proper anesthesia throughout the surgery. A local anesthetic (Lurocaine®, 100 μL at 0.67 mg/kg) was applied at the location of the scalp incision before the procedure. A median incision revealed the skull which was drilled at the level of the targeted structure. At the end of the surgery, an analgesic solution of buprenorphine (Buprecare®, 1 mg/kg) was injected subcutaneously to prepare awakening, and supplemented the following recovery days if necessary.

### Tetrode recordings

Mice were deeply anaesthetized with i.p. injection of chloral hydrate (8%, 400 mg/kg), supplemented as required to maintain optimal anesthesia throughout the experiment. After placement of the animal in the stereotaxic apparatus, the saphenous vein was catheterized (30G needle fitted into polyethylene tubing PE10 connected to a Hamilton syringe) for i.v. administration of either saline or nicotine. The scalp was opened and a hole was drilled in the skull above the location of the targeted structure (IPN and/or VTA). Experiments were carried out with a Thomas Recording Mini-Matrix fitted with independently movable tetrodes (Tip shape A, Z = 1-2 MΩ) to record neuronal activity, and a stainless-steel cannula (OD 120 μm) for MPEG_4_Ch and dye injections. Each recording consisted in a baseline period of 5 minutes before injection of either saline or nicotine, selected pseudo-randomly. Subsequent injections were spaced by at least 5 minutes after saline and 15 minutes after nicotine. For nicotine dose-response experiments, mice received two to four injections of nicotine at doses of 7.5, 15, 30 and/or 60 μg/kg (pseudo-randomly administered).

Electrophysiological signals were acquired with a 20-channel pre-amplifier included in the Mini Matrix connected to an amplifier (Digital Lynx SX 32 channels, Neuralynx) digitized and recorded using Cheetah software (Neuralynx). Spikes were detected (CSC Spike Extractor software, Neuralynx) and sorted using a classical principal component analysis associated with a cluster cutting method (SpikeSort3D software, Neuralynx). The electrophysiological activity was sampled in the VTA (coordinates: 3.0 to 3.4 mm posterior to bregma, 0.3 to 0.6 mm lateral to midline, and 4 to 4.8 mm below brain surface) and in the IPN (coordinates: 3.3 to 3.6 mm posterior to bregma, 0.35 to 0.5 mm lateral to midline, and 4.2 to 4.9 mm below brain surface). We used a 5° angle for both IPN and simultaneous IPN plus VTA recordings.

Firing frequency was measured on successive windows of 60 s shifted by 15-s time steps (45 s overlapping period). Basal neuronal activity was defined on at least five-minute recording. For each neuron, the firing frequency was rescaled as a percentage of its baseline value averaged during 3 minutes before i.v. injection. The responses to both saline and nicotine are thus presented as a percentage of variation from baseline (mean ± S.E.M.). To classify neurons as “activated” or “inhibited”, we performed the following analysis. First, we calculated the maximal variation from the baseline per neuron, within the first 3 minutes following nicotine injection. Neurons displaying an increase in firing frequency were defined as activated, while neurons displaying a decrease in firing frequency were defined as inhibited. For the dose-response curves, neurons were classified as activated or inhibited based on their response to the highest nicotine dose received. For saline injections, the polarity of the variation was defined based on the response to nicotine (i.e., in activated neurons, we considered that saline also increased activity). We used a bootstrapping method to identify neurons that significantly responded to nicotine injections. Baseline spike intervals were randomly shuffled 1000 times. For each neuron we determined the percentile from the shuffled data corresponding to the nicotine-evoked response (maximum or minimum frequency after nicotine injection). Neurons were individually considered as responsive to nicotine injection if this percentile was ≥ 0.98 (activated) or ≤ 0.02 (inhibited). The effect of nicotine was assessed by comparing the maximum of firing frequency variation induced by nicotine and saline injection. For activated (respectively inhibited) neurons, the maximal (respectively minimal) value of firing frequency was measured within the response period (3 minutes) that followed nicotine or saline injection. To assess the proper location of recordings, injections of dye (Chicago Sky Blue 6B, Sigma-Aldrich) were performed at the mean stereotaxic coordinates of recordings (500 nL at 10 nL/s). Recordings in mice where dye location was outside of the targeted structure were excluded from the analysis. Spontaneously active pDA neurons were identified on the basis of previously established electrophysiological criteria (regular firing rate; firing frequency between 1 and 10 Hz) and proper *post hoc* dye location in the VTA. For chemogenetic experiments, MPEG_4_Ch was infused (250 μM in aCSF at the rate of 10 nL/s) within either the VTA (500 nL) or the IPN (700 nL) at least an hour before recordings started. All the data were analyzed with R (https://www.r-project.org).

### Fiber photometry

8-week-old WT mice were injected with AAV1-syn-jGCaMP7c (pGP-AAV-syn-jGCaMP7c variant 1513-WPRE, titer 1.1e^13^ vg/mL, Addgene, MA, USA, https://www.addgene.org/105321/) in either the VTA (AP −3.10 mm, ML ± 0.5 mm, DV −4.5 mm from bregma, 1μL, 100 nL/min) or the IPN (AP −3.50 mm, ML ± 0.40 mm, DV − 4.7 mm from bregma, 700 nL, 100 nL/min). Optical fibers (200 μm core, NA = 0.39, Thor Labs) coupled to a stainless-steel ferule (1.25 mm diameter) were implanted after virus injection slightly above (0.05 to 0.2 mm) those coordinates, and fixed to the skull with dental cement (SuperBond, Sun medical). Recordings started at least 4 weeks after surgery.

Fluorescent levels were recorded using a Doric Lenses 1-site 2-color fiber photometry system. The fiber photometry console was connected to the LED driver to control two connectorized LED in lock-in mode (CLEDs, 405 nm CLED modulated at 333.786 Hz and 465 nm CLED modulated at 220.537 Hz) that were connected to their respective ports on the Mini Cube through an optic patch cord. Light stimulation and recorded fluorescence were transmitted through an optical fiber (400 μm core, NA = 0.39, Thorlabs) connected both to the animal’s implanted optical fiber via a zirconia sleeve and to the sample port on the Mini Cube. Finally, the photoreceiver converting recorded light to electrical signals (AC Low setting, New Focus 2151 Visible Femtowatt Photoreceiver, New Focus, San Jose, CA, USA) was connected to the Mini Cube through an optic path cord (600 μm core, NA = 0.48, FC-FC, Doric Lenses) fitted on a fiber optic adapter (Doric Lenses) and to the fiber photometry console. Signal was acquired through Doric Neuroscience Studio software (version 5.2.2.5) with a sampling rate of 12.0 kS/s (kilosamples per second) and a low-pass filter with a cutoff frequency of 12.0 Hz.

We assessed changes in GCaMP activity in response to i.v. injections of saline or nicotine. Mice were deeply anesthetized with an i.p. injection of chloral hydrate (8%, 400 mg/kg), supplemented as required to maintain optimal anesthesia throughout the experiment. Intravenous administration of saline or nicotine was carried out through a catheter (30G needle connected to polyethylene tubing PE10) connected to a Hamilton syringe, into the saphenous vein of the animal. The injection protocol was the same as in tetrode recordings.

All fiber photometry data were analyzed on R software. First, data were down-sampled by a 100-factor. We subtracted the mean value of “autofluorescence” (signal acquired after each recording with the same parameters, but without the optic fiber attached to the mouse) to the signal. We then fitted an exponential to this signal and subtracted it before adding an offset equal to the mean of the signal before detrending to account for the slow decay of the signal due to bleaching during recording. We defined a baseline fluorescence value (F0) as the mean fluorescence of the signal during 120 s before injection time, for each injection (saline and nicotine) individually. We then calculated normalized variation in fluorescence (ΔF/F) as (F-F0)/F0 for each injection. The analysis was carried out by averaging each ΔF/F obtained for each condition (all saline or nicotine injections done in IPN implanted (n =7) mice, same for saline or nicotine in VTA (n = 5) mice) and mean data were smoothed using a normal kernel fit (bandwidth = 120). For each injection (saline and nicotine), maximum fluorescence was detected within a 180 s window after injection.

### Conditioned place preference (CPP)

The conditioned place preference (CPP) experiments were performed in a Y-maze apparatus (Imetronic, Pessac, France) with one closed arm and two other arms with manually operated doors. Two rectangular chambers (11 x 25 cm) with different cues (texture and color) are separated by a center triangular compartment (side of 11 cm) used as a neutral compartment. One pairing compartment has grey textured floor and walls and the other one has smooth black and white striped walls and floor. The CPP apparatus was illuminated at 100 lux during the experiments. All animals were handled for at least five days before starting the experiment. The first day (pretest) of the experiment, without previous habituation to the apparatus, mice (n = 6-8 animals/group) explored the environment for 900 s (15 min) and the time spent in each compartment was recorded. Conditioning was unbiased: pretest data were used to segregate the animals with equal bias so each group has an initial preference score almost null, indicating no preference on average. On day 2, 3 and 4, animals received an i.p. injection of saline (0.9% NaCl) or nicotine tartrate (0.2 mg/kg, in saline), and were immediately confined to one of the pairing chambers for 1200 s (20 min). Groups were balanced so that animals did not all receive nicotine in the same chamber. On the same day, at least 5 hours after the first pairing, mice received an injection of the alternate solution (nicotine or saline) and were placed in the opposite pairing chamber. On day 5 (test), animals were allowed to freely explore the whole apparatus for 900 s (15 min), and the time spent in each chamber was recorded. The preference score is expressed in seconds and is calculated by subtracting pretest from test data. Trajectories and time spent on each side are calculated based upon animal detection. Place preference and locomotor activity were recorded using a video camera, connected to a video-track system. A home-made software (Labview 2014, National Instruments) tracked the animal, recorded its trajectory (30 frames per s) for the duration of each session.

For chemogenetic experiments, MPEG_4_Ch (250 μM in aCSF, 700 nL at 100 nL/min) or an equivalent volume of aCSF was injected under light gas anesthesia 3 days before the start of CPP paradigm. The experiments were performed in a double-blind fashion. Behavioral data were collected using home-made LabVIEW (National Instruments) and analyzed using ezTrack^45^.

### Open field and elevated o-maze tests

The open field test was run one week before the o-maze test, on the same batch of animals (n = 7-8/group). All animals were handled for at least five days before starting the open field test. The open field consisted of a circular arena (74 cm diameter), illuminated at 100 lux. Each mouse could move freely in the apparatus for 15 minutes. Total distance traveled (m), and time spent in the center area (diameter of 44 cm) were measured.

The elevated O-maze (EOM) apparatus consists of two open (stressful) and two enclosed (protecting) elevated arms that together form a zero or circle (diameter of 47 cm, height of 52 cm, 7 cm-wide circular platform). The EOM apparatus was illuminated at 150 lux in open arms and at 120 lux in closed arms. Mice were placed in the o-maze apparatus and recorded for 15 minutes. The time spent exploring the open arms, which indicates the anxiety level of the animal, was measured. Behavioral data was collected using a video camera, connected to a video-track system. LabVIEW (National Instruments) was used to track the animals and record their trajectory (30 frames per s) for the duration of each test. All data was analyzed using ezTrack^45^.

### Immunostaining

After euthanasia, brains were rapidly removed and fixed in 4% paraformaldehyde. After a period of at least three days of fixation at 4°C, serial 60-μm sections were cut from the midbrain with a vibratome. Immunostaining experiments were performed as follows: free-floating brain sections were incubated for 1 hour at 4°C in a blocking solution of phosphate-buffered saline (PBS) containing 3% bovine serum albumin (BSA, Sigma; A4503) and 0.2% Triton X-100, and then incubated overnight at 4°C with a chicken anti-GFP antibody (Life technologies Molecular Probes, A-6455) at 1:500 dilution, in PBS containing 1.5% BSA and 0.2% Triton X-100. The following day, sections were rinsed with PBS, and then incubated for 3 hours at 22-25°C with Alexa488-conjugated anti-chicken secondary antibodies (Jackson ImmunoResearch, 711-225-152) at 1:1000 dilution in a solution of 1.5% BSA in PBS. After three rinses in PBS, slices were wet-mounted using Prolong Gold Antifade Reagent (Invitrogen, P36930). Microscopy was carried out with an epifluorescent microscope (Leica), and images captured using a camera and analyzed with ImageJ.

### Statistical analysis

All statistical analyses were carried out using the R software with home-made routines. No statistical methods were used to predetermine sample sizes. In all figures, data are plotted as mean ± SEM. Total number (n) of observations in each group and statistics used are indicated in figure and/or figure legend. Unless otherwise stated, comparisons between means were carried out using parametric tests (Student’s t-test, one-way or two-way repeated measures ANOVA) when parameters followed a normal distribution (Shapiro-Wilk normality test with p > 0.05), and non-parametric tests (here, Wilcoxon or Mann-Whitney U-test) when this was not the case. Homogeneity of variances was tested preliminarily and the t-tests were Welch-corrected if needed. Likewise, one-sample comparisons to 0 were carried out using a t-test when the parameter followed a normal distribution and with a Wilcoxon test otherwise. Multiple comparisons were Holm-Bonferroni corrected. Comparison between the cumulative distributions was carried out using a Kolmogorov-Smirnov test (p > 0.05 was considered to be not statistically significant). Proportion of nicotine-activated VTA neurons in sham versus MPEG_4_Ch experiments (Figure 3E) were compared with Pearson’s Chi-squared test.

## Supplementary figures

**Figure S1:**
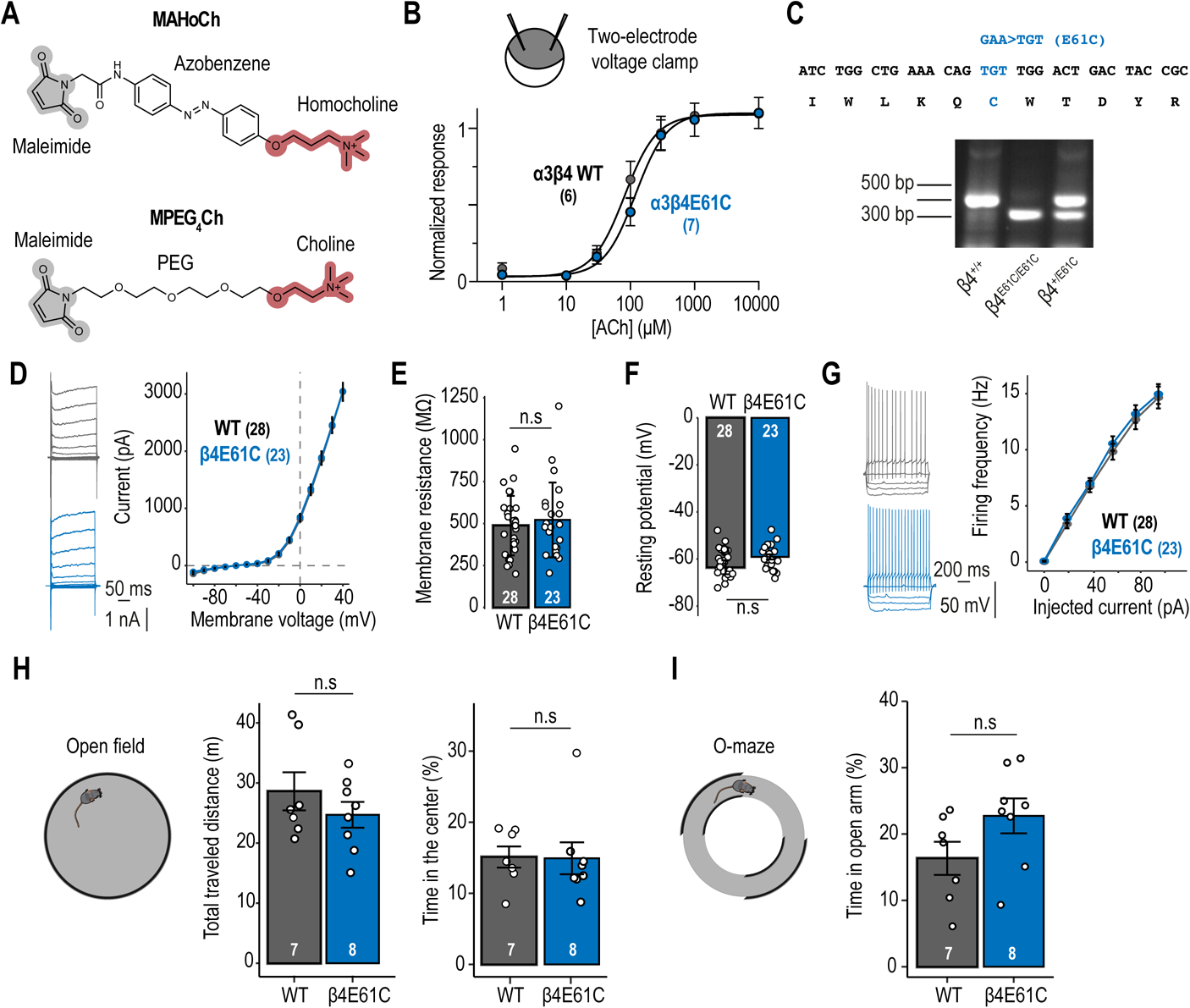
New chemogenetic tools for nAChR control. **A.** Chemical structures of the photoswitchable tethered ligand MAHoCh (top)^30^ and the suicide antagonist MPEG_4_Ch (bottom). **B.** Bottom, acetylcholine dose-response curve for α3β4 (EC_50_ = 82 μM, n_H_ = 1.7) and α3β4E61C (EC_50_ = 120 μM, n_H_ = 1.7) recorded in Xenopus oocytes. **C.** Top, glutamate at position 61 in the β4 nAChR subunit was mutated to cysteine (E61C). Bottom, PCR to genotype the β4E61C mutant mouse. **D.** Representative voltage-gated potassium currents (left) and current-voltage relationship (right) recorded from IPN neurons of WT and β4E61C mice. **E.** Membrane resistance of IPN neurons from WT and β4E61C mice. **F.** Resting membrane potential of IPN neurons from WT and β4E61C mice. **G.** Representative (left) and average (right) firing frequency vs. current relationships recorded from IPN neurons of WT and β4E61C mice. **H.** Open field test. Total distance traveled and time spent in the center for WT and β4E61C mice. **I.** Time spent in the open arm of the elevated O-maze for WT and β4E61C mice.

**Figure S2:**
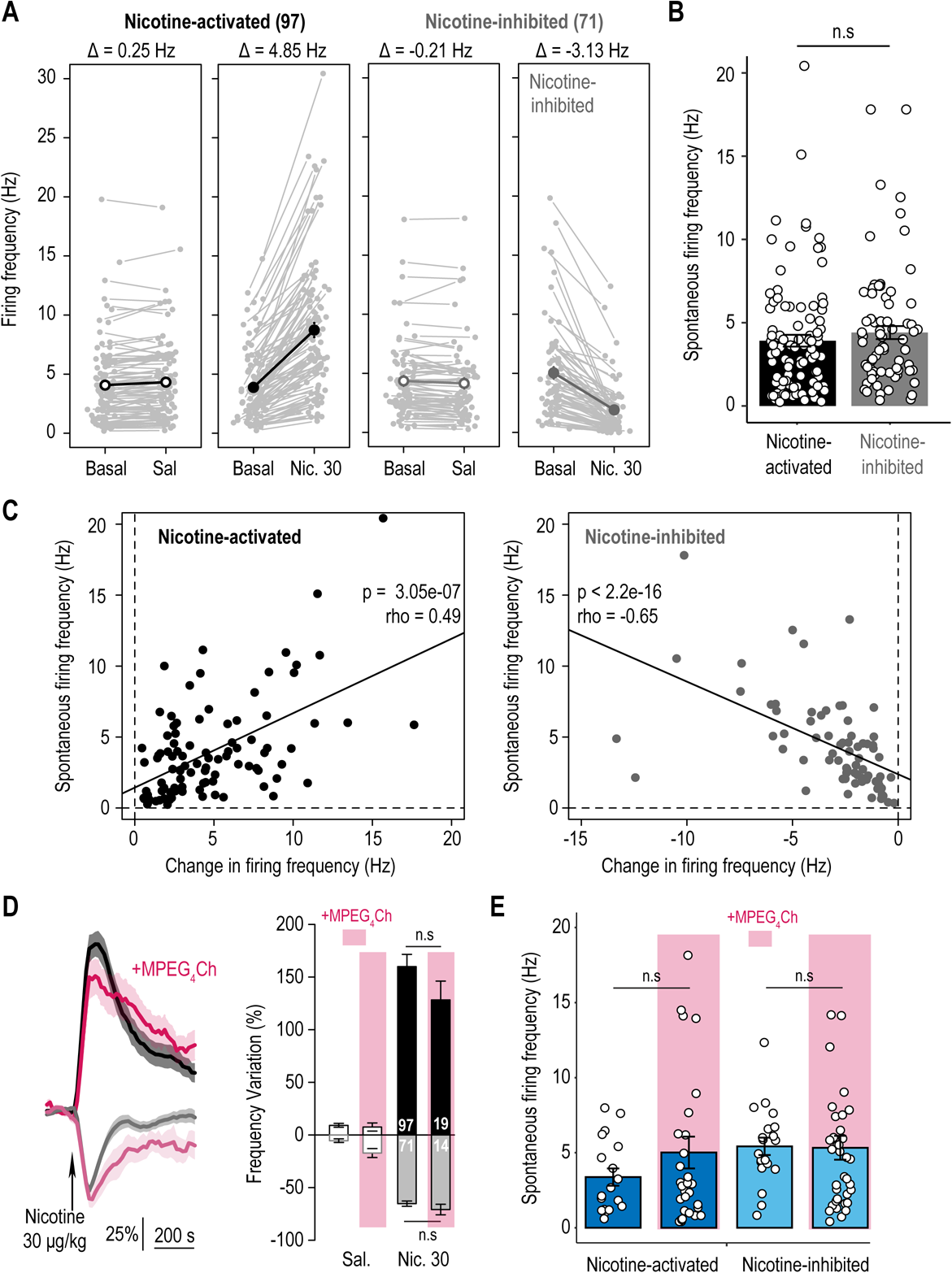
Electrophysiological characterization of the response of IPN neurons to nicotine injection. **A.** Firing rate under basal conditions and after saline of nicotine injections, for nicotine-activated and -inhibited IPN neurons. **B.** Basal firing frequency of nicotine-activated and -inhibited IPN neurons. **C.** Correlation between the basal firing frequency and the change in firing frequency following an injection of nicotine, for nicotine-activated and inhibited neurons. **D.** Time course (left) and firing frequency variation (right) after i.v. nicotine (or saline) injections, for nicotine-activated and -inhibited IPN neurons of WT mice treated or not with MPEG_4_Ch in the IPN. **E.** Spontaneous activity of nicotine-activated and -inhibited IPN neurons of βE61C mice, with and without MPEG_4_Ch treatment.

**Figure S3:**
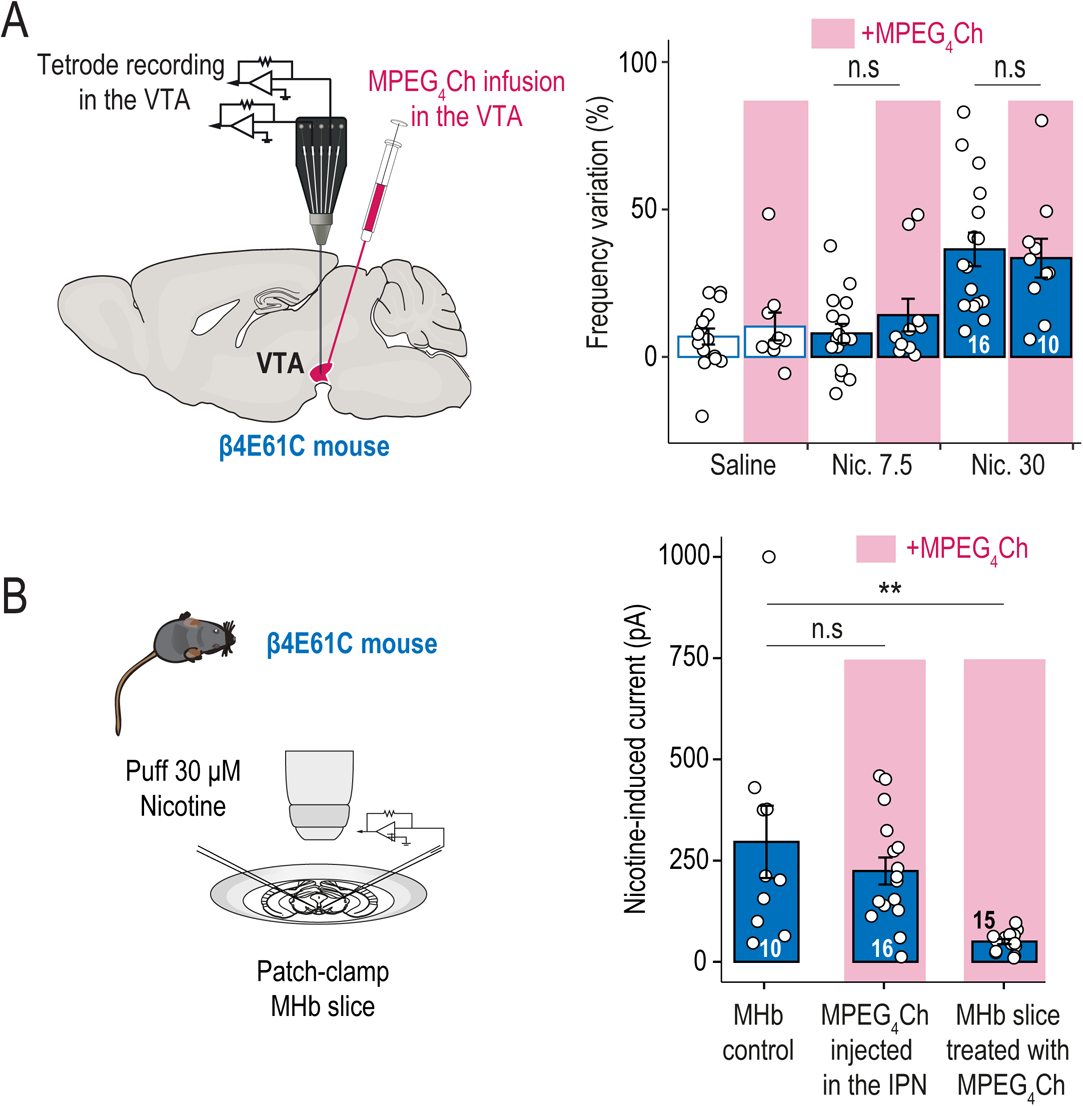
Effect of MPEG_4_Ch in other brain structures. **A.** Left, tetrode recording in the VTA of β4E61C mice, after infusion of MPEG_4_Ch in the VTA. Right, firing rate variation from baseline induced by nicotine or saline i.v. injection, in VTA neurons of β4E61C mice infused with MPEG_4_Ch in the VTA. β4E61C mice infused with ACSF (Sham) in the IPN were used as controls (same as in Figure 3E). **B.** Left, patch-clamp recording in MHb slices from β4E61C mice. Right, average current induced by a puff of nicotine (30 μM) for MHb control slices, compared with that recorded from MHb slices of β4E61C mice infused with MPEG_4_Ch in the IPN, and with that recorded from MHb slices directly treated with MPEG_4_Ch.

